# A non-canonical JAGGED1 signal to JAK2 mediates osteoblast commitment in cranial neural crest cells

**DOI:** 10.1101/421305

**Authors:** Archana Kamalakar, Melissa S. Oh, Yvonne C. Stephenson, Samir A. Ballestas-Naissir, Michael E. Davis, Nick J. Willett, Hicham M. Drissi, Steven L. Goudy

## Abstract

During craniofacial development, cranial neural crest (CNC) cells migrate into the developing face and form bone through intramembranous ossification. Loss of JAGGED1 (JAG1) signaling in the CNC cells is associated with maxillary hypoplasia or maxillary bone deficiency (MBD) in mice and recapitulates the MBD seen in humans with Alagille syndrome. JAGGED1, a membrane-bound NOTCH ligand, is required for normal craniofacial development, and *Jagged1* mutations in humans are known to cause Alagille Syndrome, which is associated with cardiac, biliary, and bone phenotypes and these children experience increased bony fractures. Previously, we demonstrated deficient maxillary osteogenesis in *Wnt1*-*cre;Jagged1^f/f^* (Jag1CKO) mice by conditional deletion of *Jagged1* in maxillary CNC cells. In this study, we investigated the JAG1 signaling pathways in a CNC cell line. Treatment with JAG1 induced osteoblast differentiation and maturation markers, *Runx2* and *Ocn*, respectively, Alkaline Phosphatase (ALP) production, as well as classic NOTCH1 targets, *Hes1* and *Hey1.* While JAG1-induced *Hes1* and *Hey1* expression levels were predictably decreased after DAPT (NOTCH inhibitor) treatment, JAG1-induced *Runx2* and *Ocn* levels were surprisingly constant in the presence of DAPT, indicating that JAG1 effects in the CNC cells are independent of the canonical NOTCH pathway. JAG1 treatment of CNC cells increased Janus Kinase 2 (JAK2) phosphorylation, which was refractory to DAPT treatment, highlighting the importance of the non-canonical NOTCH pathway during CNC cells osteoblast commitment. Pharmacologic inhibition of JAK2 phosphorylation, with and without DAPT treatment, upon JAG1 induction reduced ALP production and, *Runx2* and *Ocn* gene expression. Collectively, these data suggest that JAK2 is an essential component downstream of a non-canonical JAG1-NOTCH1 pathway through which JAG1 stimulates expression of osteoblast-specific gene targets in CNC cells that contribute to osteoblast differentiation and bone mineralization.

## 1. Introduction

Craniofacial defects like maxillary hypoplasia occur due to aberrant craniofacial development, which in humans occurs within the first 8 week of embryonic development (1). During normal craniofacial process, the first brachial arch develops in a pair of mandibular processes and a pair of maxillary processes, predominantly made up of cranial neural crest (CNC) cells that arise from the neural placode (2, 3). The maxillary processes continue to elongate laterally as palate shelves, starting at the primitive stomodeum adjacent to the tongue. The palate shelves then elevate above the tongue, fuse in the midline, and become the hard and soft palate. Intramembranous ossification of the anterior palate occurs following CNC cells osteoblast differentiation, giving rise to the hard palate, which gives shape to the maxillary bone and structure to the midface (4-6).

JAGGED1 (JAG1) is a cell surface ligand that signals through cell-cell contact via the NOTCH pathway and orchestrates maxillary development in mice by controlling proliferation, extracellular matrix production, osteoblast commitment, and vascular branching (7, 8). The interaction between the membrane-bound JAG1 and membrane-bound NOTCH receptor causes the proteolytic cleavage, by enzymes including the crucial γ-secretase, of the NOTCH intracellular domain (NICD), which then translocates to the nucleus and forms a complex with RBP-Jκ, to commence crucial gene expression of classic NOTCH targets including *Hes1* and *Hey1* (9, 10). *Jag1* is required early in craniofacial development and, its global deletion leads to embryonic death at E9.5 due to cranial hemorrhage (11). *Jag1* mutations in humans are associated with Alagille syndrome, which is associated with cardiac, biliary, facial (maxillary hypoplasia) and bony phenotypes (12-15). Our laboratory recently described that conditional deletion of *Jag1* altered CNC cells migration and differentiation, creating a mouse model of maxillary hypoplasia, a phenocopy of Alagille syndrome (7). Pups that result from conditional knockout of *Jag1* die of starvation at P21 and display maxillary bone hypoplasia (7). Thus, JAG1 signaling is required for normal maxillary development.

JAG1 is known to signal through the canonical NOTCH pathway wherein, the binding of the JAG1 ligand to one of the NOTCH receptors (1-4) on an adjacent cell, leads to cleavage of the NOTCH receptor intracellular domain (NICD), which then gets translocated into the nucleus to control gene transcription of *Hes1* and *Hey1* (16, 17). In this study, we determined the contributions by which JAG1 directs neural crest cells to osteoblast commitment via a JAG1-NOTCH non-canonical pathway. In this study, we confirmed prior reports of JAG1-NOTCH canonical osteoblast induction and described the role of JAG1-dependent non-canonical signaling through the JAK2 pathway during CNC cells osteoblast commitment.

## 2. Methods

### 2.1 Cell culture

A mouse cranial neural crest cell line, O9-1 cells (18) (Millipore sigma, SCC049) were seeded on a layer of Matrigel (Fisher, CB-40234) (1:50 dilution in 1X PBS). The cells were maintained and passaged using alpha-MEM (Gibco, 12571071) + 10% FBS (Atlanta biologics,S11150) + 1% antibiotics (Penicillin/Streptomycin) (Gibco, 15240-062) (Figure: S1A). ***Osteogenic cultures:*** The capacity of O9-1 cells to mineralize the surrounding matrix was tested by providing osteogenic media (OM): α-MEM containing 10% FBS, 1% penicillin/streptomycin, 100μg/mL ascorbic acid (Fisher, A62-500), 5 mM β-glycerophosphate (Sigma, G9422-50G) and 100ng/mL BMP2 (Creative Biomart, BMP2-01H). To assure that osteogenic media was essential for matrix mineralization, control wells were incubated in growth media (GM): α-MEM containing 10% FBS and 1% penicillin/streptomycin. Half-feeds were given every 2 days.

### 2.2 JAGGED1 immobilization

Dynabeads Protein G (19) (Invitrogen 10003D) were first resuspended in the vial. 50 μL (1.5 mg) of Dynabeads were transferred to a tube, where the beads were separated from the solution using a magnet. Recombinant JAG1-Fc (10μg) (Creative biomart, JAG1-3138H) and Fc fragment (10μg) (Abcam, ab108557) alone diluted in 200 μL PBS with Tween-20 (Fisher, BP337-500) were added to the Dynabeads. The beads + proteins were incubated at 4°C with rotation for 16 hours. Thereafter the tubes were placed on the magnet and the supernatant was removed. Then the tubes were removed from the magnet and the beads-Ab complex was resuspended in 200 μL PBS with Tween-20 to wash by gentle pipetting. The wash buffer was also separated from the beads-Ab complex using the magnet. A final suspension of the beads in media was used as treatment (Figure S1).

### 2.3 Western blots

O9-1 cells were seeded on Matrigel in 6-well plates (Costar, 3516) and cultured until 100% confluency. The cells were then serum starved for 16 hours prior to treatments. The cells were first treated with 5μM WP1066 (inhibitor of JAK2 phosphorylation) (Millipore, S65784-10mg) and 15μM DAPT (a γ-secretase inhibitor) (Sigma, D5942-25mg) for 1 hours prior to stimulus with: JAG1-Fc-Beads complex (10μg), Fc-Beads complex (10μg), BMP2 (100ng/ml), 200μM Etoposide (activator of JAK2 phosphorylation) (Sigma, E1383-25mg), 5μM WP1066 and 15μM DAPT. Cells were then lysed using RIPA buffer (Thermoscientific, 89900) containing a protease inhibitor (Roche, 05892791001) and a phosphatase inhibitor (Roche, 04906845001) to obtain whole cell proteins that were resolved on an 8% SDS-PAGE gel and then transferred to a nitrocellulose membrane. The blots were then placed in 1.5% BSA (Sigma, A2153-100G) + 0.1% Tween-20 for 30 minutes and thereafter incubated at 4°C for 16 hours with a primary antibody against phosphorylated JAK2 (Cell signaling, 8082S) and later total-JAK2 (Cell signaling, 3230S) at 1:1000 dilution in 1.5% BSA + 0.1% Tween-20. The blots were moved to secondary antibody (Cell signaling, 7074P2) prepared at a 1:3000 dilution in 1.5% BSA + 0.1% Tween-20 and incubated at room temperature for 1 hour, then washed using TBS (Amresco, 0307-5L) + 0.1% Tween-20 3 times, 5 minutes each wash. Blots were developed by chemiluminescence using the Bio-rad Chemidoc MP imaging system available at the Emory Children’s Center Pediatric research equipment core. ***PCR:*** For gene expression, we used qPCR, as previously described (20). Total RNA was isolated using the TRIzol reagent (Ambion, 15596018) according to the manufacturer’s protocol. cDNA was generated from 1μg total RNA using the High capacity cDNA reverse transcriptase kit (Applied biosystems, 4368813).

**Supplementary Figure S1:**
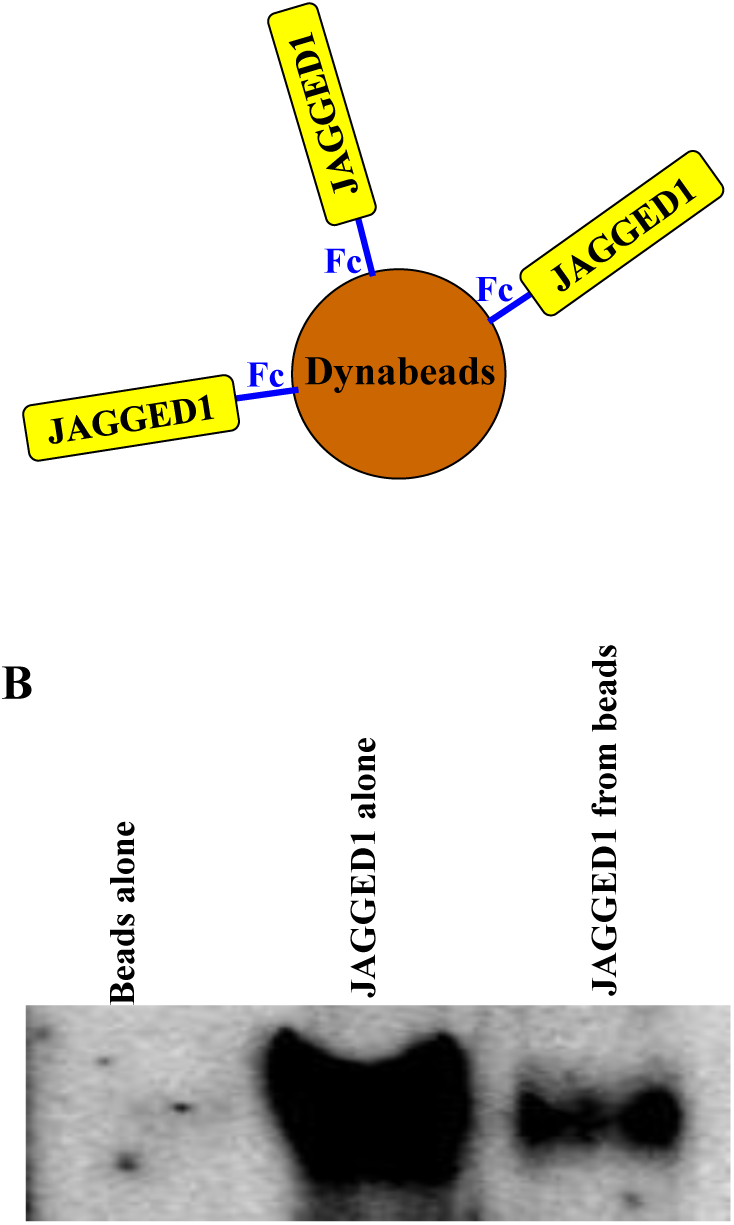
**Recombinant JAGGED1-Fc protein was immobilized on Protein G coated Dynabeads,** as shown in (A). (B) Immobilization success was determined by eluting the JAGGED1 attached to the Dynabeads and probed for by immunoblotting.

Real-time PCR analysis was done with iQ SYBR green supermix (Bio-Rad, 1708882) in the Bio-Rad iCycler for 40 cycles. Primer pairs are shown in Supplemental Table 1. The expression levels are calculated using the ΔΔCT method. The threshold cycle (CT) represents the PCR cycle at which an increase of the reporter fluorescence above the baseline is first detected. The fold change in expression levels, R, is calculated as follows: R=2–ΔΔCT (where R = 2 (ΔCT treated-ΔCT control)). The abundance of all transcripts was normalized to the level of GAPDH RNA expression.

### 2.4 Alkaline Phosphatase (ALP) assay

O9-1 cells were cultured for 10 days in osteogenic media in 24-well plates; we measured ALP activity of the cells as a marker of osteoblast committment. The cells were fixed using 4% PFA (Sigma, F8775) for 10 minutes at 4°C and then gently rinsed using tap water in a small water container. The plates were allowed to dry for 10 minutes, thereafter, p-Nitrophenyl Phosphate Liquid Substrate System Liquid (Sigma, N7653-100) is presented to the cells as substrate for ALP and incubated at 37°C for 30 minutes. The solution turns yellow indicating hydrolysis of p-Nitrophenyl Phosphate by ALP. The absorbance of this product was read by a spectrophotometer at 405nm.

### 2.5 Crystal Violet assay

In order to obtain quantitative information about the density of adherent cells in a multi well plate, a Crystal Violet assay was performed. The ALP assay product was carefully removed from the wells by washing with tap water gently in a small water container, and allowed to dry for 10 minutes. A 0.2% crystal violet (VWR, 0528-500G) solution prepared in a 2% ethanol (Acros organic, 61509-0020) in water solution was added to the cells and incubated for 1 minute at room temperature. The excess crystal violet was washed using tap water by immersion in a small water container. To quantify the crystal violet taken up by the cells, the stain was solubilized in a 1:10 dilution of acetic acid in water, and the absorbance was read at 570nm.

### 2.6 Statistics

Data were analyzed by analysis of variance (ANOVA) with Tukey’s post-test. All data are presented as mean +/- SD. p<0.05 between groups was considered significant and are reported as such.

**Supplementary Table 1:**
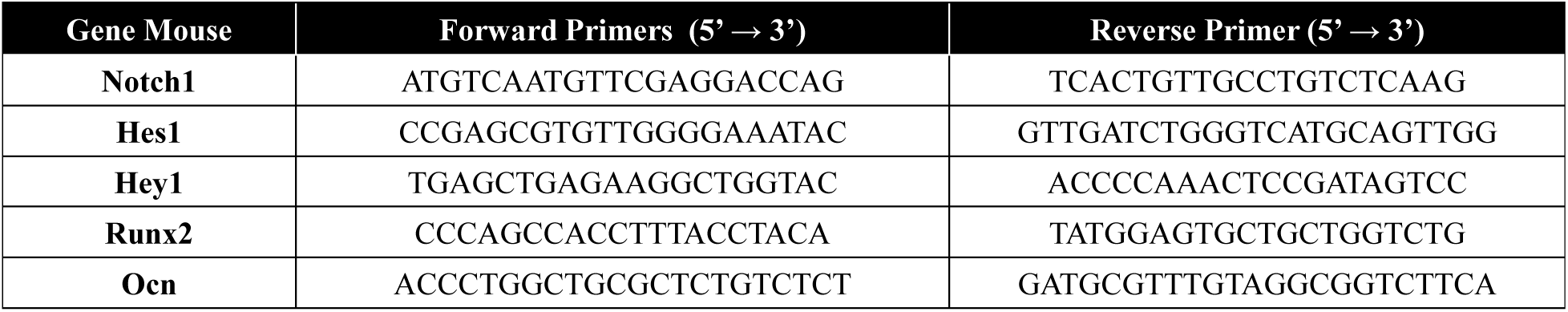
Primers used for PCR analysis. Primers were designed using the NCBI-NIH Primer-BLAST tool for all genes.

## 3. Results

### 3.1 JAGGED1 upregulates Notch1 receptor expression as well as expression of NOTCH1 targets Hes1 and Hey1

O9-1 cells (Figure: S2A) express the NOTCH1 receptor (Figure: S2B) and are known to mineralize in the presence of osteogenic media as shown in Supplemental Figure: 2C. O9-1 cells were treated with osteogenic media (OM) alone, dynabeads alone and Fc fragment-beads complex (10μg) as controls or JAG1-Fc-Beads complex (10μg) as shown in Figure 1. BMP2 (100ng/ml) is a known activator of the NOTCH canonical pathway (21) and was thus used as a positive control. A γ-secretase inhibitor, DAPT (15μM) was used as a NOTCH canonical pathway inhibitor as it inhibits the cleavage of the NOTCH intracellular domain (NICD). We observed significant increase in *Notch1* gene expression within 3hrs of JAG1 treatment compared to untreated (OM alone) and BMP2-treated cells (Figure 1A). As expected, treatment with DAPT, a γ-secretase inhibitor of NOTCH canonical pathway, ± JAG1, resulted in increased *Notch*1 gene expression, suggesting a compensatory mechanism in response to lack of NOTCH activity. We also observed that JAG1 induced classic NOTCH1 targets, *Hes1* and *Hey1*, expression levels were predictably decreased after DAPT (NOTCH inhibitor) treatment (Figure 1B & 1C). This indicates that JAG1 activates and functions through the NOTCH canonical pathway in O9-1 cells.

**Figure 1:**
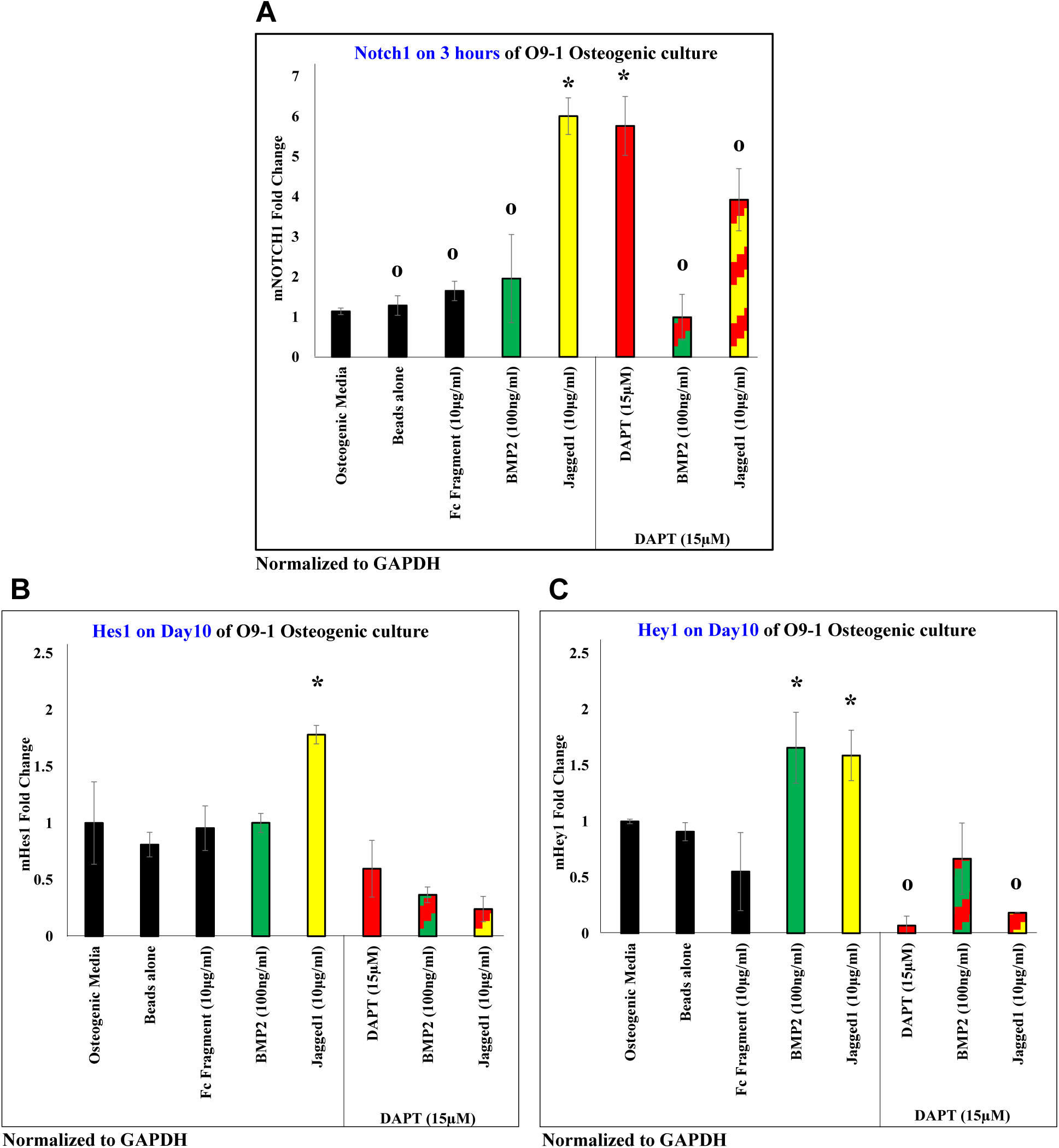
**JAGGED1 induces the NOTCH pathway,** by inducing expression of *Notch1* receptor in (A), and expression of classic NOTCH targets (B) *Hes1* and (C) *Hey1*. The expression of all genes induced by JAG1 was observed to be inhibited by a canonical NOTCH pathway inhibitor, DAPT (red bar, red patterned bars). Dynabeads bound recombinant JAG1-Fc fragment (yellow) or BMP2 (green, positive control) were used. Growth media, osteogenic media, dynabeads alone and dynabeads bound Fc-fragment were used as controls (black bars). (n=3) (Similar symbols = no difference, different symbols = significant difference) (p<0.05)

### 3.2 JAGGED1 induces the differentiation of neural crest cells into osteoblasts

Previously we published that loss of *Jag1* results in maxillary bone hypoplasia (7), and here we demonstrate the direct induction of osteoblastogenesis by JAG1 in O9-1 cells. Alkaline phosphatase (ALP) assay on JAG1-induced O9-1 cells, demonstrated that ALP activity was significantly increased with JAG1 treatment (Figure 2A). Expression of osteoblast differentiation and maturation markers, *Runx2* and *Ocn*, respectively, were increased with JAG1 treatment, indicating that JAG1 can independently induce osteoblast commitment (Figure 2B & 2C). JAG1 induced similar levels of *Runx2* and *Ocn* gene expression with and without DAPT treatment (Figure 2B & 2C), indicating that the JAG1 effects in the O9-1 cells are largely independent of NICD cleavage, i.e. the NOTCH canonical pathway. Increased expression of *Runx2* following DAPT treatment was observed and has been identified by other authors, however the expression of *Ocn*, a late osteoblast marker, was not increased in the DAPT treated cells, suggesting that blockage of NICD cleavage alone is not sufficient to induce osteoblast differentiation. In BMP2 treated cells, addition of DAPT reduced the expression of *Runx2* and *Ocn* demonstrating that BMP2 signals, in part, through the canonical NOTCH pathway, a finding that has been described by other investigators.

**Supplementary Figure S2:**
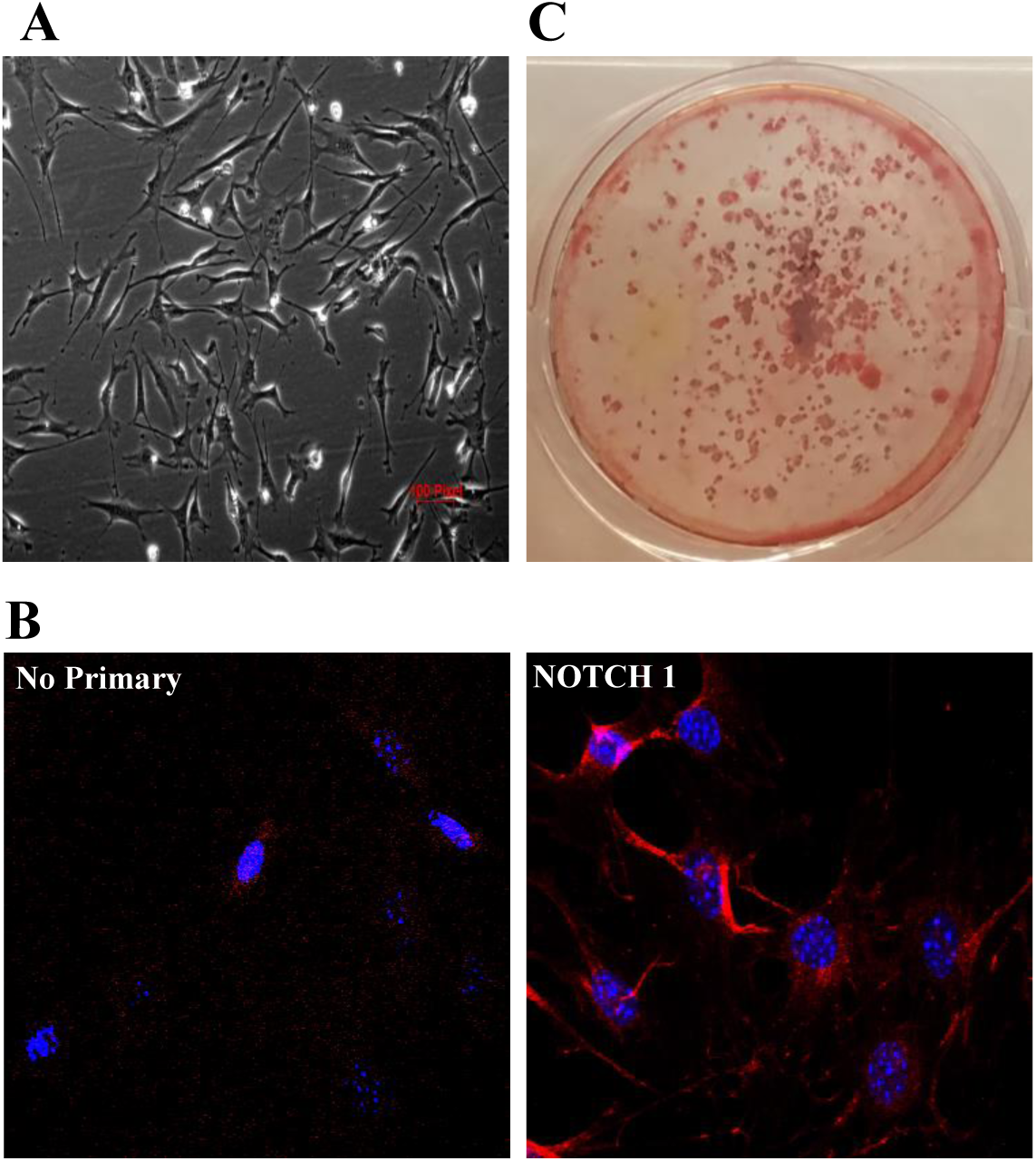
**Characterization of O9-1 cells** as seen in culture in (A) express the NOTCH1 receptor (Red staining) demonstrated by an Immunofluorescent staining assay in (B). (C) O9-1 cells were driven towards osteoblastogenesis in the presence of β-Glycerophosphate and Ascorbic acid until day 28 cells when they were stained with Alizarin Red to indicate that these cells are capable of inducing mineralization.

**Figure 2:**
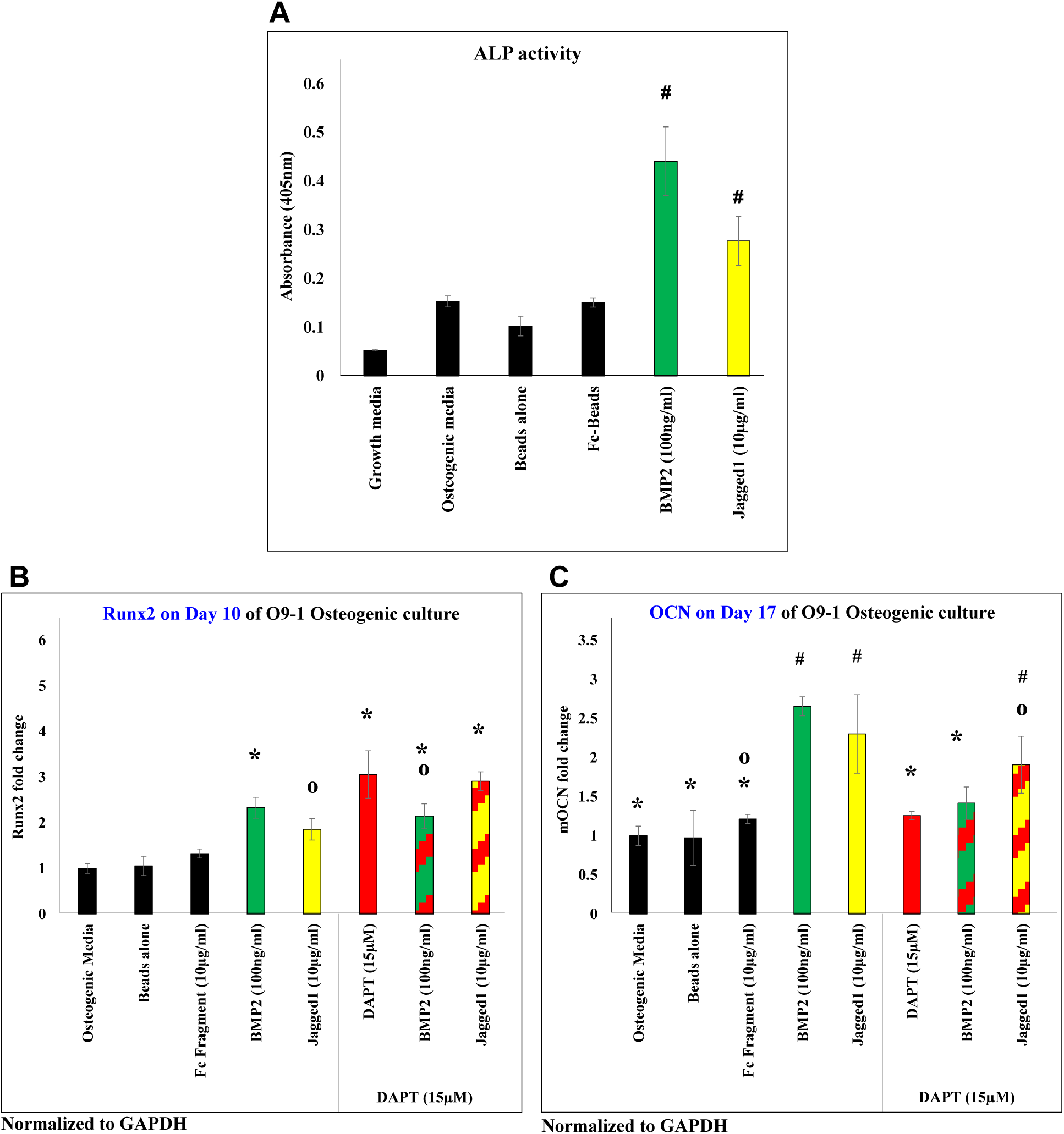
**JAGGED1 induces osteoblastogenesis,** as demonstrated by measuring the (A) alkaline phosphatase activity of O9-1 cells when treated with dynabeads bound recombinant JAG1-Fc fragment (yellow) or BMP2 (green, positive control). Growth media, osteogenic media, dynabeads alone and dynabeads bound Fc-fragment were used as controls (black bars). The canonical NOTCH pathway inhibitor DAPT (red bar, red patterned bars) did not inhibit JAG1-induced gene expression of (B) early osteoblast marker, *Runx2*, (C) late osteoblast marker, *Ocn*. (n=3) (Similar symbols = no difference, different symbols = significant difference) (p<0.05)

### 3.3 JAGGED1 non-canonical signaling during CNC cells osteoblast commitment via JAK2 phosphorylation

To elucidate the non-canonical pathways activated downstream of JAG1 we examined known contributors to osteoblast commitment including p38, ERK, SMAD159, AKT, JAK2 (22-26). SMAD159 and AKT phosphorylation was not induced by JAG1 (Figures S3A) whereas, the phosphorylation of ERK and P38 was observed to be constitutively active in O9-1 cells (Figures S3B). JAG1 treatment of O9-1 cells was associated with increased JAK2 phosphorylation (Figure 3A) and this phosphorylation was observed to be refractory to DAPT treatment (Figure 3B), demonstrating the importance of JAG1-NOTCH non-canonical pathway signaling during CNC cells osteoblast commitment. Pharmacological inhibition of JAK2 phosphorylation using WP1066 (5μM) after JAG1-stimulation decreased the *Notch1* (Figure 4A). The expression levels of *Hes1* and *Hey1* (Figure 4B & 4C), remained unchanged, as they are classic canonical NOTCH pathway targets and this pathway was stimulated by JAG1. Although, the *Runx2* and *Ocn* gene expression (Figure 5A, & 5B), were decreased along with *Notch1* (Figure 4A), suggesting that JAK2 phosphorylation is an essential component downstream of the non-canonical JAG1-NOTCH1 pathway during osteoblast commitment. Further, pharmacologic inhibition of JAK2 phosphorylation ± DAPT upon JAG1 induction synergistically reduced ALP production (Figure 5C). Collectively, these data suggest that JAK2 is an essential component downstream of a non-canonical JAG1-NOTCH1 pathway through which JAG1 stimulates expression of osteoblast-specific gene targets that contribute to osteoblast differentiation.

**Supplementary Figure S3:**
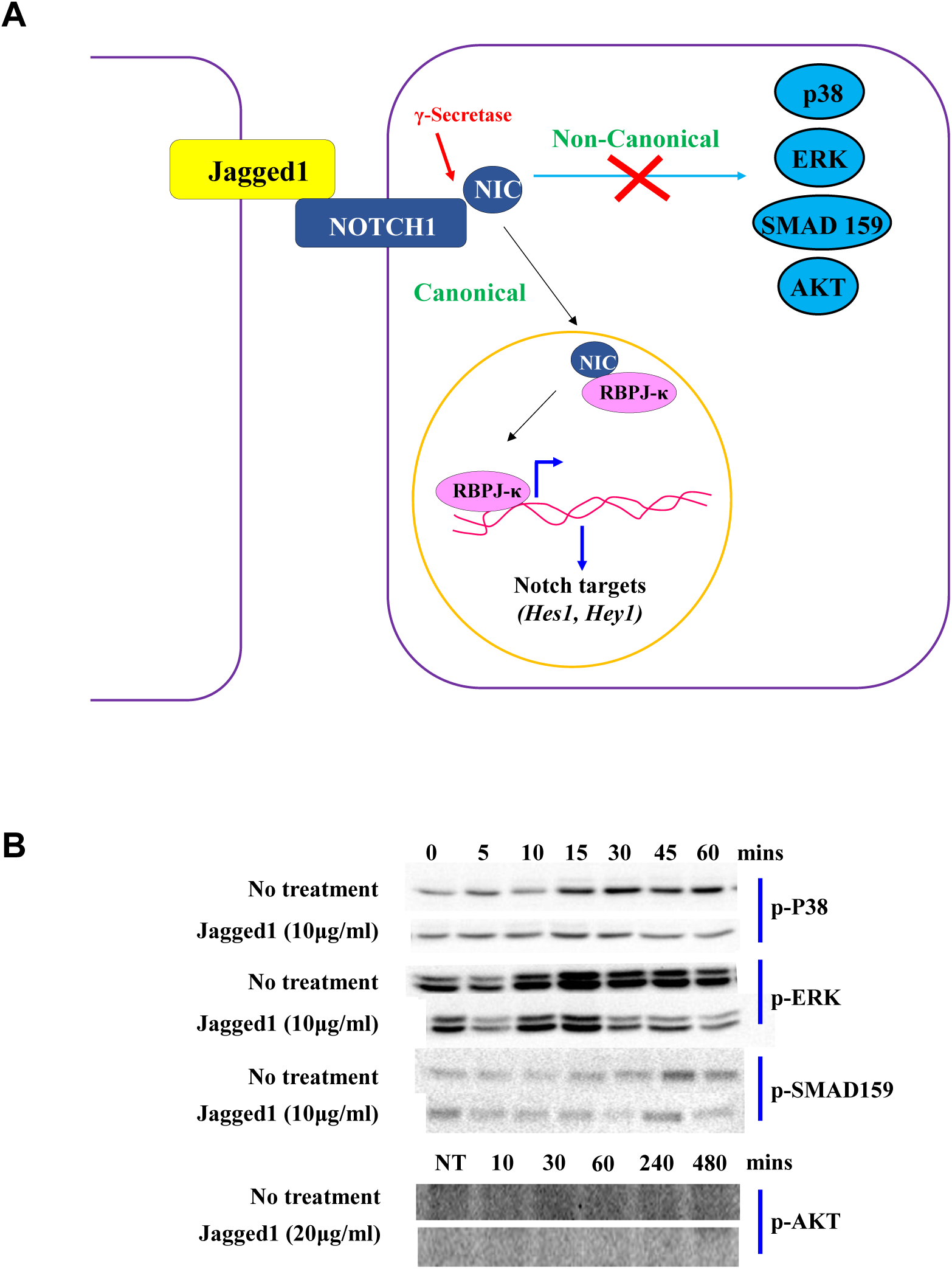
**(A) Illustration of the hypothesis** that JAGGED1 activates a non-canonical pathway downstream of the NOTCH1 receptor. Although p38, ERK, AKT, SMAD159 are not phosphorylated downstream of JAGGED1-NOTCH1 compared to No treatment controls as shown in (B).

**Figure 3:**
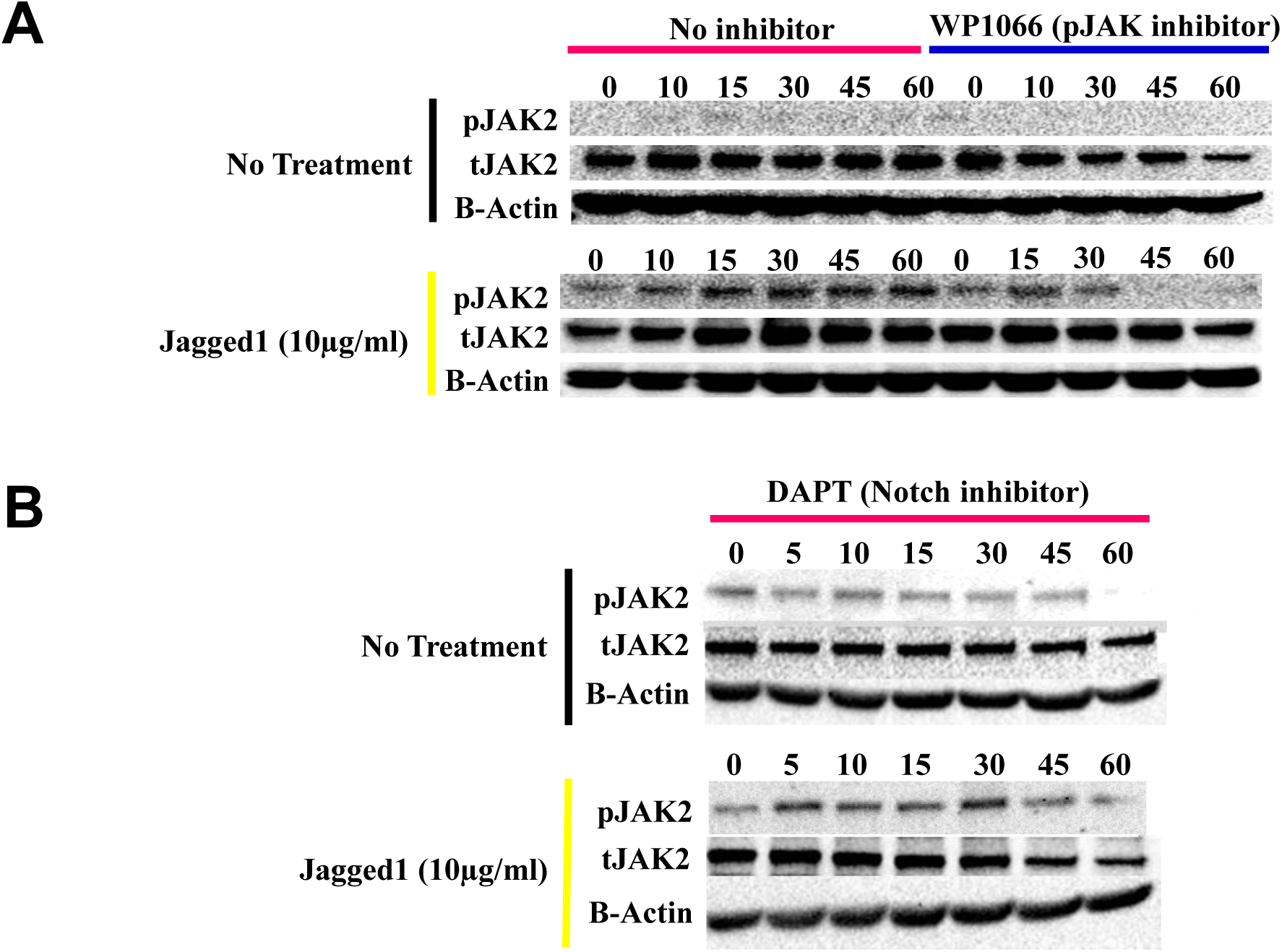
JAGGED1 induces a non-canonical NOTCH pathway. (A) Lysates obtained from O9-1 cells treated with dynabeads bound Fc-fragment, BMP2 and etoposide, both known inducers of JAK2 phosphorylation (data not shown) and dynabeads bound recombinant JAG1-Fc fragment, were probed for phosphorylated JAK2. (B) O9-1 cells were treated with a NOTCH canonical pathway inhibitor, DAPT, prior to adding treatments. Lysates were then probed for JAK2 phosphorylation.

**Figure 4:**
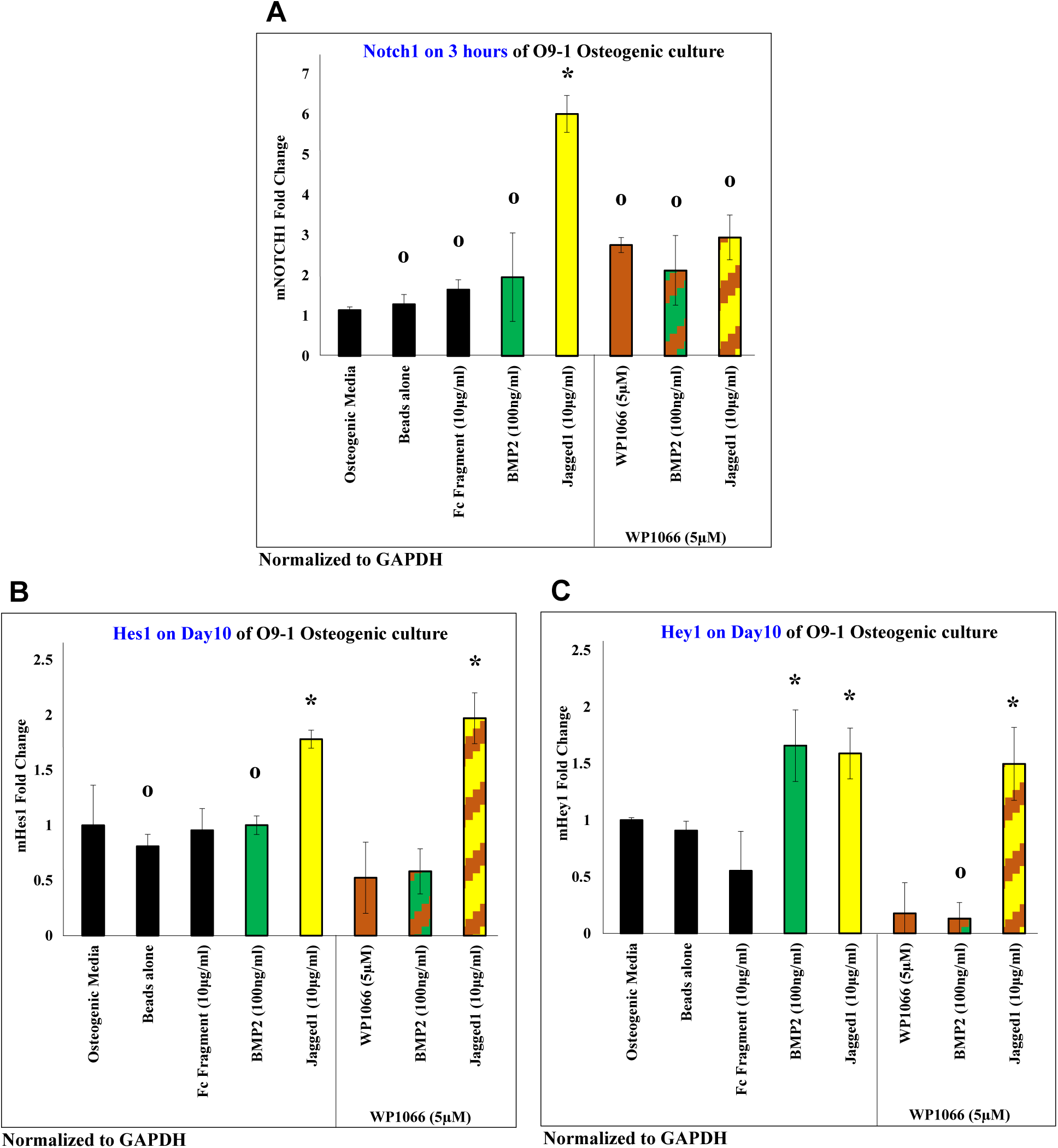
**JAGGED1 induces the NOTCH canonical pathway**, demonstrated by treatment of O9-1 cells with an inhibitor of JAK2 phosphorylation, WP1066 (brown bar) prior to treatments with dynabeads bound recombinant JAG1-Fc fragment (yellow-brown patterned bar) or BMP2 (green-brown patterned bar) *Notch1* receptor expression in (A) was significantly decreased, although expression of classic NOTCH targets (B) *Hes1* and (C) *Hey1* were uninhibited in the presence of WP1066. (n=3) (Similar symbols = no difference, different symbols = significant difference) (p<0.05)

**Figure 5:**
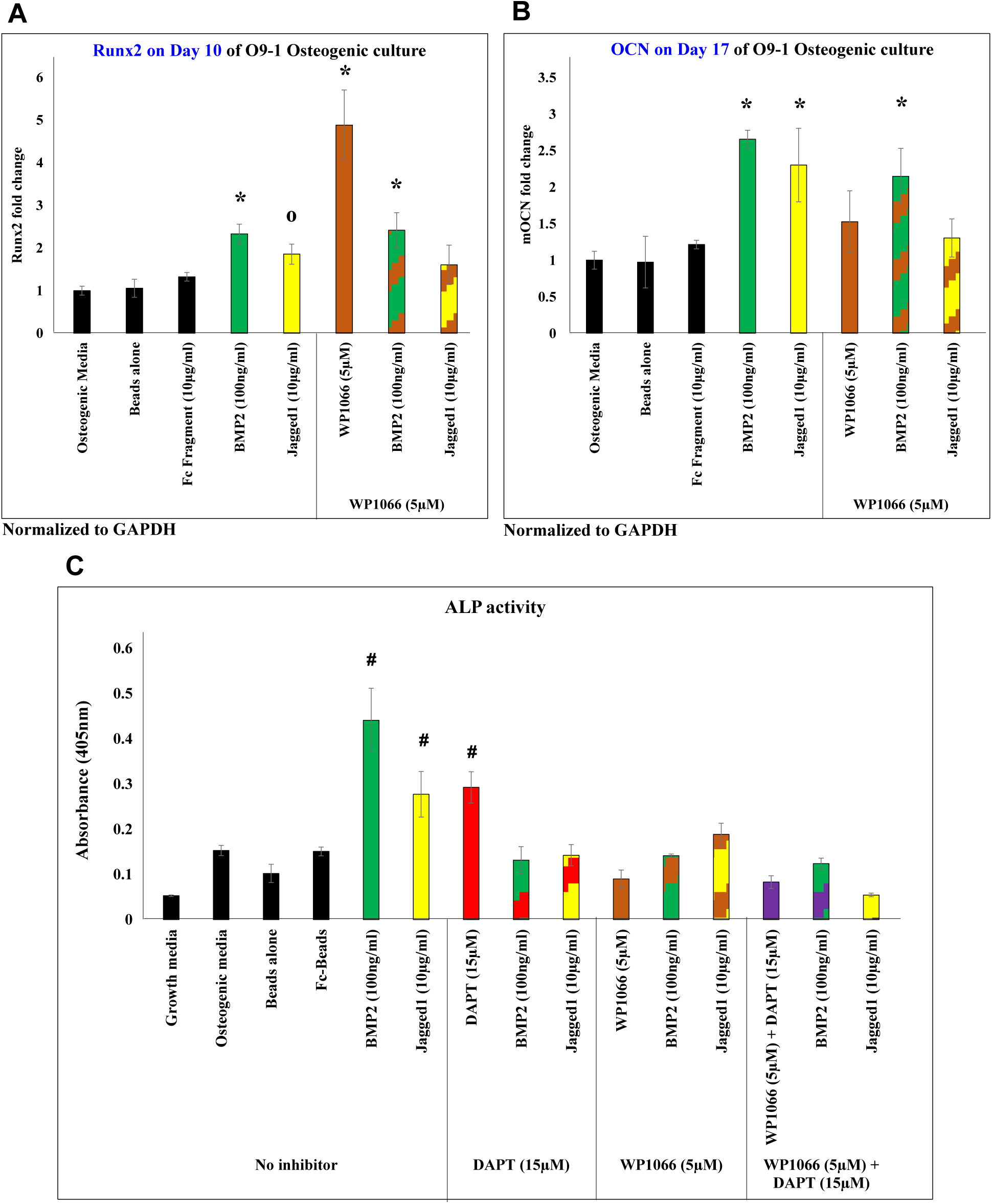
**JAGGED1 induces osteoblastogenesis via NOTCH non canonical signal through phosphorylated JAK2**, as demonstrated by measuring the gene expression of (A) early osteoblast marker, *Runx2*, (B) late osteoblast marker, *Ocn*, (C) alkaline phosphatase (ALP) activity of O9-1 cells treated with an inhibitor of JAK2 phosphorylation prior to treatment with dynabeads bound recombinant JAG1-Fc fragment (yellow-brown patterned bar) or BMP2 (green-brown patterned bar). *Runx2* and *Ocn* were significantly decreased in JAG1 + WP1066-treated cells. ALP activity was significantly decreased in JAG1 + DAPT-treated (yellow-red patterned bar) as well as JAG1 + WP1066-treated O9-1 cells but it was synergistically decreased in O9-1 cells treated with JAG1 along with both DAPT and WP1066 (yellow-purple patterned bar). (n=3) (Similar symbols = no difference, different symbols = significant difference) (p<0.05).

## 4. Summary & Discussion

JAG1 is a membrane-bound NOTCH ligand required for normal craniofacial development, and *JAG1* mutations in humans are known to cause maxillary bone hypoplasia (13, 27). Alagille syndrome, the consequence of a *Jag1* or *Notch2* mutation, is associated with a mélange of symptoms including pathological bone phenotypes such as, reduced bone mineral density and a higher propensity to bony fractures (28). The canonical NOTCH pathway has been implicated in a plethora of biological processes including, embryonic development, craniofacial development, bone homeostasis, endothelial cell fate, angiogenesis and T-cell lineage commitment, when specifically activated by the ligand JAG1 (27, 29-31). The role of NOTCH is known to be dimorphic, (32) wherein *Notch*’s pathological gain of function leads to osteosclerosis and *Notch*’s loss of function leads to increase in osteoclastogenesis, and subsequent bone loss. Conditional loss of *Jag1* (Jag1CKO) in CNC cells was associated with reduced proliferation and osteoblast commitment, leading to maxillary hypoplasia, starvation and death of pups at post-natal day 21 (7, 13). Evaluation *ex vivo* cultures of the Jag1CKO mouse CNC cells demonstrated decreased mineralization, suggesting that the JAG1-NOTCH axis is essential for the maturation of neural crest cells into osteoblasts (7). However, it remains unclear how the JAG1-NOTCH pathway controls differentiation of CNC cells during intramembranous ossification.

JAG1 is a definitive NOTCH1 ligand and induces the NOTCH1 canonical pathway, which others and we have demonstrated by examining JAG1-induced expression of classic NOTCH1 target, *Hes1* and *Hey1* (Figure 1B & 1C) (33). We and others have also demonstrated, as in Figure 2B & 2C, that JAG1 induces early and late osteoblast markers, *Runx2* and *Ocn*, respectively, along with inducing ALP activity, *in vitro*, in O9-1 cells, independent of BMP2 (Figure: 2A, 2B & 2C) suggesting that the JAG1-NOTCH1 axis is essential during commitment of neural crest cells into the osteoblast lineage (33, 34). Inhibition of the canonical JAG1-NOTCH1 pathway, using a γ-secretase inhibitor, DAPT, reduced ALP activity demonstrating that JAG1 can induce CNC cells towards osteoblast commitment and maturation via the canonical NOTCH pathway. However, DAPT treatment did not alter the gene expression levels of the JAG1-induced *Runx2* and *Ocn*, which suggests that JAG1 signals through canonical and non-canonical NOTCH pathways during CNC cells osteoblast commitment.

The canonical NOTCH pathway has been shown to cross talk with various other non-canonical signaling pathway molecules including MAPK (ERK, P38), Src, NF-κB and PI3K (22, 35-37). In addition, the ERK, P38, SMAD, AKT and JAK2 pathways have been, frequently shown to regulate osteoblast differentiation (22-26). Prior reports have demonstrated that constitutively active NOTCH, *in vitro*, inhibits BMP2-induced bone formation whereas, transiently active NOTCH enhances bone formation (36). Dahlqvist et, al demonstrated that the NICD binds to BMP2-induced SMAD3 to facilitate induction of bone formation. JAG1-activated NOTCH pathway has been described to regulate the maintenance of the osteoprogenitor pool, thereby suppressing osteoblast differentiation (38). The contrasting data about the bone inhibitory and inductive functions of JAG1-NOTCH1 are likely related to the origin of cell and the situational context depending on which JAG1 signals. Examination of the non-canonical NOTCH signaling pathways has received less attention but this may help explain the contradictory functions of JAG1-NOTCH1 in the literature. In Supplemental figure 3, we evaluated the activation of numerous non-canonical osteogenic pathways including ERK, P38, SMAD159, AKT and JAK2 following JAG1 treatment of CNC cells. JAG1 did not activate ERK, P38, SMAD159 or AKT, but we successfully demonstrated that JAG1 phosphorylates JAK2 (Figure: 3).

JAK2 is a signaling molecule required during osteogenesis and skeletogenesis, and complete abrogation of JAK2 signaling is associated with embryonic death at E12.5 (39). JAK2 is phosphorylated downstream of both, Polycystin, a key player in mechanotransduction during skeletogenesis (40) and Growth hormone (GH), a regulator of bone growth and metabolism (41). BMP2 and dexamethasone are known to synergistically activate JAK2 to result in increased levels of ALP indicating bone formation (42). Pharmacologically inhibited JAK2 phosphorylation in JAG1 treated CNC cells demonstrated that the JAK2 inhibitor, WP1066 was able to suppress the expression of *Runx2* and *Ocn* as well as the ALP activity. Inhibition of JAK2 phosphorylation and the NOTCH canonical pathway, using WP1066 and DAPT, respectively, demonstrated that the ALP activity was significantly and synergistically decreased further corroborating that a JAG1 signal to JAK2 mediates osteoblast commitment in CNC cells. During metastasis of pancreatic ductal adenocarcinoma, the canonical NOTCH pathway itself has been shown to cross talk with the JAK/STAT pathway, wherein, HES proteins facilitate JAK2 and STAT3 to form a complex thereby promoting STAT3 phosphorylation and activation (43), thus, suggesting that JAG1-JAK2 signaling may occur via the canonical as well as the non-canonical NOTCH pathway in osteoblast commitment of CNC cells.

## 5. Conclusions

In this study, we introduce a novel, non-canonical JAG1-JAK2 signaling pathway to induce osteoblast commitment. We report that JAG1 can independently activate JAK2 in CNC cells, downstream of NOTCH1, which further induces the expression of osteoblast markers, *Runx2* and *Ocn*, along with increasing ALP activity, indicating a non-canonical signal that drives CNC cells to commit to the osteoblast lineage. In order to understand thoroughly the preferred mode of action by which JAG1 leads CNC cells to osteoblast commitment in the context of the cell’s microenvironment and effectors, it is exigent to evaluate the cell context (Figure 6A), stimulators (Figure 6B), upstream and downstream mediators (Figure 6C & 6D) of the JAG1-induced JAK2 signaling, and its interactions with the NOTCH canonical pathway (Figure 6E). Currently, there are no FDA approved therapies for JAG1-associated bone loss but our identification of JAG1’s mode of action and its ability to independently induce bone formation could change the paradigm of treatment for maxillary hypoplasia in Alagille Syndrome patients.

**Figure 6:**
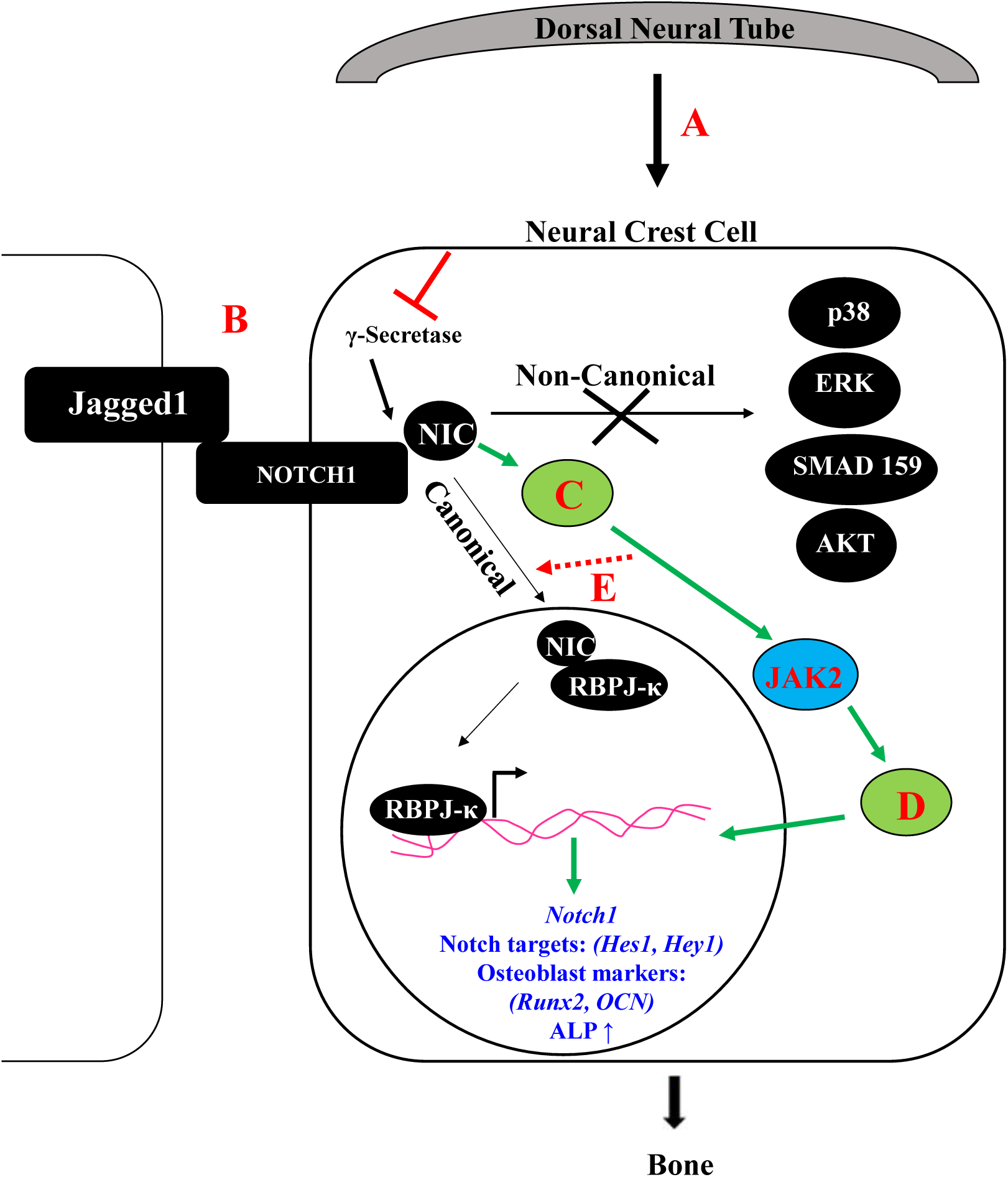
Illustration of mechanism of JAGGED1-induced osteoblast commitment of neural crest cells. JAG1 induces osteoblast formation not only by activating the canonical NOTCH pathway but also by activating a non-canonical NOTCH signal involving the phosphorylation of JAK2. Future directions include understanding (A) the role of the origin of the neural crest cells during intramembranous ossification, (B) action of JAG1 through other NOTCH receptors (2-4), (C) upstream and (D) downstream targets of JAK2 as well as whether there can be (E) crosstalk between the canonical and non-canonical pathways during JAGGED1 induced osteoblast formation.

## 6. Acknowledgements

Study design: AK, MO, MD, NW, HD and SG. Study conduct: AK. Data collection: AK, MO, YS and SB. Data analysis: AK, MO, YS and SB. Data interpretation: All authors. Drafting manuscript: AK and SG. Revising manuscript content: SG, MD, NW and HD. Approving final version of manuscript: All authors. AK and SG take responsibility for the integrity of the data analysis. The Oral Maxillofacial Surgery Foundation funded this project (Funding ID: 2591).

## Abbreviations

CNC: Cranial Neural Crest
JAG1: JAGGED1
Jag1CKO: JAGGED1 Conditional Knockout
JAK2: Janus Kinase
ALP: Alkaline Phosphatase

## 7. References

1. Wilderman A, VanOudenhove J, Kron J, Noonan JP, Cotney J. High-Resolution Epigenomic Atlas of Human Embryonic Craniofacial Development. Cell Rep. 2018;23(5):1581-97.

2. Wu T, Chen G, Tian F, Liu HX. Contribution of cranial neural crest cells to mouse skull development. Int J Dev Biol. 2017;61(8-9):495-503.

3. Meulemans D, Bronner-Fraser M. Gene-regulatory interactions in neural crest evolution and development. Dev Cell. 2004;7(3):291-9.

4. Welsh IC, Hagge-Greenberg A, O’Brien TP. A dosage-dependent role for Spry2 in growth and patterning during palate development. Mech Dev. 2007;124(9-10):746-61.

5. Ferguson MW. Palate development. Development. 1988;103 Suppl:41-60.

6. Franz-Odendaal TA. Induction and patterning of intramembranous bone. Front Biosci (Landmark Ed). 2011;16:2734-46.

7. Hill CR, Yuasa M, Schoenecker J, Goudy SL. Jagged1 is essential for osteoblast development during maxillary ossification. Bone. 2014;62:10-21.

8. Benedito R, Roca C, Sorensen I, Adams S, Gossler A, Fruttiger M, et al. The notch ligands Dll4 and Jagged1 have opposing effects on angiogenesis. Cell. 2009;137(6):1124-35.

9. Hori K, Sen A, Artavanis-Tsakonas S. Notch signaling at a glance. J Cell Sci. 2013;126(Pt 10):2135-40.

10. Wilson JJ, Kovall RA. Crystal structure of the CSL-Notch-Mastermind ternary complex bound to DNA. Cell. 2006;124(5):985-96.

11. Xue Y, Gao X, Lindsell CE, Norton CR, Chang B, Hicks C, et al. Embryonic lethality and vascular defects in mice lacking the Notch ligand Jagged1. Hum Mol Genet. 1999;8(5):723-30.

12. Oda T, Elkahloun AG, Pike BL, Okajima K, Krantz ID, Genin A, et al. Mutations in the human Jagged1 gene are responsible for Alagille syndrome. Nat Genet. 1997;16(3):235-42.

13. Humphreys R, Zheng W, Prince LS, Qu X, Brown C, Loomes K, et al. Cranial neural crest ablation of Jagged1 recapitulates the craniofacial phenotype of Alagille syndrome patients. Hum Mol Genet. 2012;21(6):1374-83.

14. Ziesenitz VC, Loukanov T, Glaser C, Gorenflo M. Variable expression of Alagille syndrome in a family with a new JAG1 gene mutation. Cardiol Young. 2016;26(1):164-7.

15. Li L, Dong J, Wang X, Guo H, Wang H, Zhao J, et al. JAG1 Mutation Spectrum and Origin in Chinese Children with Clinical Features of Alagille Syndrome. PLoS One. 2015;10(6):e0130355.

16. Lindsell CE, Shawber CJ, Boulter J, Weinmaster G. Jagged: a mammalian ligand that activates Notch1. Cell. 1995;80(6):909-17.

17. Shimizu K, Chiba S, Saito T, Kumano K, Takahashi T, Hirai H. Manic fringe and lunatic fringe modify different sites of the Notch2 extracellular region, resulting in different signaling modulation. J Biol Chem. 2001;276(28):25753-8.

18. Ishii M, Arias AC, Liu L, Chen YB, Bronner ME, Maxson RE. A stable cranial neural crest cell line from mouse. Stem Cells Dev. 2012;21(17):3069-80.

19. Ndong JC, Stephenson Y, Davis ME, Garcia AJ, Goudy S. Controlled JAGGED1 delivery induces human embryonic palate mesenchymal cells to form osteoblasts. J Biomed Mater Res A. 2018;106(2):552-60.

20. Hill CR, Sanchez NS, Love JD, Arrieta JA, Hong CC, Brown CB, et al. BMP2 signals loss of epithelial character in epicardial cells but requires the Type III TGFbeta receptor to promote invasion. Cell Signal. 2012;24(5):1012-22.

21. Vilain N, Tsai-Pflugfelder M, Benoit A, Gasser SM, Leroy D. Modulation of drug sensitivity in yeast cells by the ATP-binding domain of human DNA topoisomerase IIalpha. Nucleic Acids Res. 2003;31(19):5714-22.

22. Rodriguez-Carballo E, Gamez B, Ventura F. p38 MAPK Signaling in Osteoblast Differentiation. Front Cell Dev Biol. 2016;4:40.

23. Ge C, Xiao G, Jiang D, Franceschi RT. Critical role of the extracellular signal-regulated kinase-MAPK pathway in osteoblast differentiation and skeletal development. J Cell Biol. 2007;176(5):709-18.

24. Fukuda T, Kohda M, Kanomata K, Nojima J, Nakamura A, Kamizono J, et al. Constitutively activated ALK2 and increased SMAD1/5 cooperatively induce bone morphogenetic protein signaling in fibrodysplasia ossificans progressiva. J Biol Chem. 2009;284(11):7149-56.

25. Choi YH, Jeong HM, Jin YH, Li H, Yeo CY, Lee KY. Akt phosphorylates and regulates the osteogenic activity of Osterix. Biochem Biophys Res Commun. 2011;411(3):637-41.

26. Darvin P, Joung YH, Yang YM. JAK2-STAT5B pathway and osteoblast differentiation. JAKSTAT. 2013;2(4):e24931.

27. Youngstrom DW, Senos R, Zondervan RL, Brodeur JD, Lints AR, Young DR, et al. Intraoperative delivery of the Notch ligand Jagged-1 regenerates appendicular and craniofacial bone defects. NPJ Regen Med. 2017;2:32.

28. Kamath BM, Loomes KM, Oakey RJ, Emerick KE, Conversano T, Spinner NB, et al. Facial features in Alagille syndrome: specific or cholestasis facies? Am J Med Genet. 2002;112(2):163-70.

29. Lehar SM, Dooley J, Farr AG, Bevan MJ. Notch ligands Delta 1 and Jagged1 transmit distinct signals to T-cell precursors. Blood. 2005;105(4):1440-7.

30. Lawal RA, Zhou X, Batey K, Hoffman CM, Georger MA, Radtke F, et al. The Notch Ligand Jagged1 Regulates the Osteoblastic Lineage by Maintaining the Osteoprogenitor Pool. J Bone Miner Res. 2017;32(6):1320-31.

31. Kume T. Ligand-dependent Notch signaling in vascular formation. Adv Exp Med Biol. 2012;727:210-22.

32. Engin F, Yao Z, Yang T, Zhou G, Bertin T, Jiang MM, et al. Dimorphic effects of Notch signaling in bone homeostasis. Nat Med. 2008;14(3):299-305.

33. Manokawinchoke J, Nattasit P, Thongngam T, Pavasant P, Tompkins KA, Egusa H, et al. Indirect immobilized Jagged1 suppresses cell cycle progression and induces odonto/osteogenic differentiation in human dental pulp cells. Sci Rep. 2017;7(1):10124.

34. Youngstrom DW, Dishowitz MI, Bales CB, Carr E, Mutyaba PL, Kozloff KM, et al. Jagged1 expression by osteoblast-lineage cells regulates trabecular bone mass and periosteal expansion in mice. Bone. 2016;91:64-74.

35. Byun MR, Kim AR, Hwang JH, Kim KM, Hwang ES, Hong JH. FGF2 stimulates osteogenic differentiation through ERK induced TAZ expression. Bone. 2014;58:72-80.

36. Nobta M, Tsukazaki T, Shibata Y, Xin C, Moriishi T, Sakano S, et al. Critical regulation of bone morphogenetic protein-induced osteoblastic differentiation by Delta1/Jagged1-activated Notch1 signaling. J Biol Chem. 2005;280(16):15842-8.

37. Hales EC, Taub JW, Matherly LH. New insights into Notch1 regulation of the PI3K-AKT-mTOR1 signaling axis: targeted therapy of gamma-secretase inhibitor resistant T-cell acute lymphoblastic leukemia. Cell Signal. 2014;26(1):149-61.

38. Blokzijl A, Dahlqvist C, Reissmann E, Falk A, Moliner A, Lendahl U, et al. Cross-talk between the Notch and TGF-beta signaling pathways mediated by interaction of the Notch intracellular domain with Smad3. J Cell Biol. 2003;163(4):723-8.

39. Li J. JAK-STAT and bone metabolism. JAKSTAT. 2013;2(3):e23930.

40. Gargalionis AN, Basdra EK, Papavassiliou AG. Polycystins and mechanotransduction in bone. Oncotarget. 2017;8(63):106159-60.

41. Joung YH, Lim EJ, Darvin P, Chung SC, Jang JW, Do Park K, et al. MSM enhances GH signaling via the Jak2/STAT5b pathway in osteoblast-like cells and osteoblast differentiation through the activation of STAT5b in MSCs. PLoS One. 2012;7(10):e47477.

42. Mikami Y, Asano M, Honda MJ, Takagi M. Bone morphogenetic protein 2 and dexamethasone synergistically increase alkaline phosphatase levels through JAK/STAT signaling in C3H10T1/2 cells. J Cell Physiol. 2010;223(1):123-33.

43. Palagani V, Bozko P, El Khatib M, Belahmer H, Giese N, Sipos B, et al. Combined inhibition of Notch and JAK/STAT is superior to monotherapies and impairs pancreatic cancer progression. Carcinogenesis. 2014;35(4):859-66.

